# The stability of lower genital tract (LGT) microbiota correlates with reproductive system function and in vitro fertilization and frozen embryo transfer outcomes in women with polycystic ovarian syndrome

**DOI:** 10.1101/2024.01.18.576212

**Authors:** Yaoyao Tu, Yixiang Zhang, Huixi Chen, Bin Wei, Yingzhou Ge, Guolian Ding, Xi Dong, Jianzhong Sheng, Li Jin, Hefeng Huang

**Affiliations:** Obstetrics and Gynecology Hospital, Institute of Reproduction and Development, Fudan University, Shanghai, 200011, China; Department of Obstetrics and Gynecology, International Peace Maternity and Child Health Hospital, Shanghai Jiao Tong University, Shanghai 200030, China; Shanghai Key Laboratory of Embryo Original Diseases, Shanghai 200030, China; Department of Obstetrics and Gynecology, the Fourth Affiliated Hospital, International Institutes of Medicine, Zhejiang University School of Medicine, Hangzhou 322000, China; Key Laboratory of Reproductive Genetics (Ministry of Education), Department of Reproductive Endocrinology, Women’s Hospital, Zhejiang University School of Medicine, Hangzhou 310058, China; Reproductive Medicine Center, Zhongshan Hospital, Fudan University, Shanghai 200032, China; CAS Center for Excellence in Molecular Plant Sciences, Shanghai Institutes for Biological Sciences (SIBS), Chinese Academy of Sciences. Shanghai 200032, China; University of Chinese Academy of Sciences, Shanghai 200032, China; Research Units of Embryo Original Diseases, Chinese Academy of Medical Sciences, Shanghai, China

**Keywords:** PCOS, lower genital tract (LGT), microbiota, *Lactobacillus spp.*, *Gardnerella spp.*, LH/FSH, menstrual cycle, FET outcomes

## Abstract

We previously found that the lower genital tract (LGT) microbiota of polycystic ovarian syndrome (PCOS) women has altered when compared to healthy reproductive-aged women, however whether this alternation will have an impact on the reproductive system function and pregnancy outcomes of PCOS patients have not yet been identified. Between 2018 and 2021, we totally enrolled 191 reproductive-aged women in three independent case-control studies, 209 vaginal and 209 cervical swabs, and 9 cervical epithelial cells samples were collected from the study subjects. Firstly, we analyzed the correlation between LGT flora and clinical characteristics of 97 women (containing 47 PCOS patients and 50 control reproductive-aged women), canonical correspondence analysis (CCA) showed that LH/FSH ratio was the most relevant factor that was related to the dominant genera in women with PCOS (r^2^=0.233, p=0.001), and menstrual cycle frequency was also strongly related to the LGT organisms (r^2^=0.116, p=0.005). Next, through analysis of 72 PCOS patients who were underwent IVF-FET therapy, the FET outcome in PNB group (PCOS with relative abundance of *Lactobacillus* ≥50%, n=57) was significant better than PDB group(PCOS patients with relative abundance of *Lactobacillus*<50%, n=15). Further, we randomly selected nine reproductive-aged PCOS patients (approximately half of them had altered LGT microbiota: *Lactobacillus spp.* ≤50%, n=5) and simultaneously collected both LGT swabs and cervical epithelial cells from them. By synchronization analysis of RNA sequencing of the cervical epithelial cells and 16S rRNA sequencing of the microbes, we found that the gene expression profiles of the cells significantly differed between the PDB (PCOS patients with relative abundance of *Lactobacillus*<50%, n=4) and PNB (PCOS with relative abundance of *Lactobacillus* ≥50%, n=5) groups. Above all, we systematically elucidate the potential relationship between LGT microbiota with the reproductive system function and IVF-FET outcomes in PCOS patients.

**Importance:** polycystic ovarian syndrome (PCOS) women always suffered from poor pregnancy outcome: high incidence of abortion, preterm-birth, and premature rupture of membranes. Although some patients have improved their reproductive outcomes through assisted reproductive technology, the risk of early clinical pregnancy loss for PCOS patients after IVF treatment still ranges from 12% to 48%. As we previously found that the LGT flora of PCOS women had significant altered when compared with healthy parallel group, and more evidence showed that the genital tract microbiota may have a critical role in the process of embryo implantation and development, here we used multiple study groups to explore the potential relationship between LGT microbiota with reproductive system function and FET outcome in PCOS patients in this study. Our findings provide a new perspective for exploring novel therapy to improve the reproductive outcomes of PCOS patients.

## Introduction

Microbiota that lives in the genital tract (GT) has aroused scientists’ interest in recent years due to their potential critical roles in women’s reproductive health (1–4). Disturbance in the composition of GT microbiota has been associated with a variety of gynecological and obstetric conditions, such as bacterial vaginitis, sexual transmitted diseases (STDs), endometritis, infertility, abortion, preterm birth, etc (5–9). Predominance implantation of *Lactobacillus spp.* can provide a low vaginal pH (∼4.0), which can maintain the immune homeostasis of the local epithelial mucosal layer. (10, 11). Considering the direct continuity of the reproductive tract, microbiota lives in the lower genital tract (LGT) might closely related to the upper genital tract (UGT) (12–14), which may influenced the UGT microenvironment and function, and further have an impact on the process of embryo implantation and development. However, so far, many studies on this subject are limited to epidemiological research, more reliable evidence are needed.

Polycystic ovary syndrome (PCOS) is the most common endocrine disease in women (affecting about 5-20% reproductive-aged women worldwide), which can cause multiple system disorders, such as reproductive function disorders (15, 16), abnormal glucose metabolism, cardiovascular diseases et al (17, 18). Reproductive-aged women with PCOS always more vulnerable to experience pregnancy loss and develop pregnancy complications, such as gestational diabetes, preeclampsia, and preterm birth, multiple risk factors have been reported to take a role in the procedure: hyperandrogenism, high LH concentration, obesity, insulin resistance and so on (17). In our previous study (19), we found that the LGT (vaginal and cervical) microbiota in PCOS women was significantly altered when compared to that of healthy reproductive-aged women: *Lactobacillus spp.* was significantly reduced in the vagina and cervical of PCOS patients, while several bacteria that are known to be related to vaginitis and adverse pregnancy outcomes, such as *Gardnerella_vaginalis_00703mash*, *Prevotella_9_other*, and *Mycoplasma hominis*, were abundant. Unlike the well-established factors that are known to affect the reproductive outcome of PCOS patients, the role of the LGT microbiota might have been underestimated (3). Also, little is known about the factors that may affect the populations of the organisms that live in the LGT of women with PCOS and whether these alterations influence local cellular function, and consequently, the reproductive outcome.

In this study, we found that the microbiota enriched in the LGT of PCOS women is positively related with higher level of luteinizing hormone/follicle-stimulating hormone (LH/FSH) ratio and prolonged menstrual cycles. When we compared the relative abundance of vaginal genera with the FET outcomes (implantation, clinical pregnancy, and live birth rates) of 72 PCOS patients who were underwent IVF-FET, we found that the *Gardnerella spp.* was significantly reduced in the live birth group (p=0.03*), while *Lactobacillus spp.* (p=0.08) was higher in the implantation group, however the difference lacks statistical significance. Through joint analysis of the16SrRNA sequencing of cervical flora and the RNA sequencing of cervical epithelial cells, we found that PCOS patients who with lower LGT abundance of *Lactobacillus spp.* (relative abundance<50%), their gene expression in the cervical epithelial cells changed remarkable when compared to PCOS patients who with higher LGT *Lactobacillus spp.* abundance. Through Gene Ontology (GO) enrichment analysis, we found that the altered biological process in cervical cells which lives with a lower abundance of *Lactobacillus spp.* (relative abundance<50%) mainly involves in the integrity of the epithelial cell barrier, cell responses to microorganisms, immune response, etc. This result shows that there is a strong correlation between flora and local epithelial function: reduced abundance of *Lactobacillus spp.* is closely related to significant epithelial cell dysfunction of PCOS patients in cervical. However, due to restricted in sampling, whether the upper genital tract epithelial cell function changed simultaneously need to be further studied.

This study provides novel insight into the LGT microbiota factors that may related with reproductive system function, and IVF-FET outcomes of women with PCOS. Our findings suggest that a regular menstrual cycle accompanied by lowered LH/FSH ratio is important for PCOS patients as it may be critical to maintain a higher abundance of *Lactobacillus spp* in LGT microbe, which might directly relate to a more healthy epithelial function and a better pregnancy outcome. Further research may consider whether we can improve the PCOS patients’ reproductive outcome by vaginal microbiota intervention.

## Materials and methods

### Sample collection and study design

In this study, 191 reproductive-aged Chinese women were recruited from the International Peace Maternity and Child Health Hospital, Obstetrics and Gynecology Hospital of Fudan University, and Zhongshan Hospital of Fudan University, between December 2018 and October 2021. All subjects were fully informed about the study objectives. PCOS was diagnosed with Rotterdam criteria, that is, presence of at least two of the following: irregular menstrual periods (oligomenorrhea or amenorrhea), polycystic ovaries, and hyperandrogenism. Patients with Cushing’s syndrome, congenital adrenal hyperplasia, thyroid disorder, hyperprolactinemia, or androgen-secreting tumors were excluded (14). Healthy women who were of similar ages as the PCOS patients were recruited as the control group. All the healthy women were attending the assisted reproductive clinic because of male-factor infertility, and their physical examination indexes were normal. Any subject with inflammation, endocrine disorder, or cancer was excluded. Women who were pregnant, nursing, or menstruating at the time of sampling were also excluded. We ascertained that the subjects had not used antibiotics within 7 days, performed cervical treatment or flushing within 5 days, or engaged in sexual activity within 48 h before the commencement of the study.

### Ethical approval

All patients provided written informed consent. The study was approved by the International Peace Maternity and Child Health Hospital (ethics approval number GKLW 2018-10), Hospital of Obstetrics and Gynecology of Fudan University (ethics approval number JIAI E2020-017), and Zhongshan Hospital of Fudan University (ethics approval number B2021-326R). PCOS patients who were undergoing in vitro fertilization (IVF)-FET were also receiving hormone replacement therapy. Vaginal and cervical swabs were collected with sterile cotton swabs, while cervical specimens were collected carefully with a vaginal dilator. The collected swabs were immediately put into a clean 2 ml DNA LoBind tube containing sterile saline on ice and transferred to a freezer at -80℃ within 2 hours. Thereafter, DNA extraction was done (19). The cervical epithelial cells were collected by inserting a brush into the cervical, rotating it clockwise 15 times, and carefully avoiding contamination from other positions. The collected brushes were put into a test tube containing 5 ml phosphate-buffered saline, placed on ice immediately, and transferred to the laboratory within 2 hours. They were then shaken at 4℃ for 5 min, centrifuged at 1,500 rpm for 5 min, washed once with phosphate-buffered saline, and centrifuged again. The supernatant was discarded, and TRIzol lysate was added to re-suspend the cells. They were then frozen with liquid nitrogen for RNA extraction and detection. All the materials used for these procedures were sterilized.

### Sequencing and analysis

#### Illumina sequencing and analysis of 16S rRNA

Total genomic DNA was extracted from each swab using the QIAGEN Power Soil Kit (12888-100, QIAGEN), following the manufacturer’s instructions (20). The bacterial 16S rRNA gene was amplified by polymerase chain reaction with the primers V3-V4. Purified amplicons were sequenced on an Illumina HiSeq 2500 platform (San Diego, CA). The acquired sequences were filtered for quality control as previously described (21), Chimera detection and removal were accomplished using the USEARCH tool in the UCHIME algorithm (22). Sequences were then split into groups according to taxonomy and assigned to operational taxonomic units (OTUs) at a 3% dissimilarity level (i.e., 97% similarity), using the UPARSE pipeline. OTUs with two or fewer sequences were removed, and to classify the representative sequences of the remaining OTUs, we used the SILVA database (release 128). In total, we obtained 18,902 OTUs, and 93.6% of them were successfully identified taxonomically.

#### RNA sequencing and analysis

To explore transcriptional differences between the two subgroups of PCOS patients and divergence in their LGT microbiota composition, we sampled cervical epithelial cells as well as LGT swabs of nine PCOS patients. The patients were divided into two groups based on the relative abundance of *Lactobacillus spp.* in the LGT. The PNB (PCOS non-depend on bacteria) group comprised patients with relative abundance of *Lactobacillus spp.* ≤50% (n=5), while the PDB (PCOS depend on bacteria) group comprised patients with relative abundance of *Lactobacillus spp.* >50% (n=4). Samples were collected and then flash-frozen in liquid nitrogen before storage at -80°C prior to RNA extraction. Total RNA was isolated with TRIzol reagent (Invitrogen, Paisley, UK). Using the Illumina HiSeq 4000 platform in the 2 × 150 bp mode, construction of cDNA libraries and subsequent sequencing were conducted by Genergy Ltd. (Shanghai, China). The statistics of the sequence data are shown as supplementary information. Each sample was independently mapped to the reference genome and subjected to expression profiling using the mode “quant” of Salmon v0.12.078, with the parameter “-validateMappings”. All independent profiles were ultimately merged to a “TPM” matrix using the “quantmerge” mode of Salmon v0.12.0. Expression profile-based principal component analysis was performed using the built-in R function “prcomp.” Highly expressed genes were selected based on the TPM rank of each treatment group, and the 500 most highly expressed genes were determined. Pathway enrichment analyses were performed with clusterProfiler (https://yulab-smu.github.io/clusterProfiler-book/). False discovery rate-adjusted multiple tests were added to the hypergeometric test. Functional enrichment analysis was performed on 497 differentially expressed genes. The results are shown in “1-differential gene (497) enrichment analysis results.” We selected the pathways from them, redrew the column chart, and screened the differential genes that were related to these pathways. We then conducted correlation analysis with the target flora, screened the relationship pairs with absolute value of the correlation coefficient >0.6, and drew the network diagram.

### Statistical Analyses

All statistical analyses were performed using R version 4.0.1. We used Shannon index to test for differences in microbiome composition among various sample groups. Since the composition of the LGT microbiota may be affected by diverse factors, we also performed canonical correspondence analysis (CCA) and Spearman correlation analysis to assess the effects of clinical variables on the microbiome composition in different groups. Results of alpha diversity tests are presented in Figures 1D and 2A, with a significance threshold of p <0.05. To test the differences in microbial community composition between the PCOS and control groups, the genus-level count data were transformed to compositional (relative) abundances to detect a high fraction of the population with sample prevalence (20%) and relative abundance (0.01%) in all samples. When bacteria with corresponding external indicators appeared in different groups, we conducted independent t-tests to compare the differences in genera under different grouping conditions (implantation, clinical pregnancy, and live birth). In Gene Ontology (GO) enrichment analysis, Fisher’s exact test was used to calculate the enrichment significance of each term in biological process, cellular component, and molecular function. Kyto Encyclopedia of Genes and Genomes (KEGG) pathway enrichment analysis was done, using hypergeometric distribution to calculate the degree of association between each path in the KEGG pathway and the differentially expressed genes.

**Figure 1.**
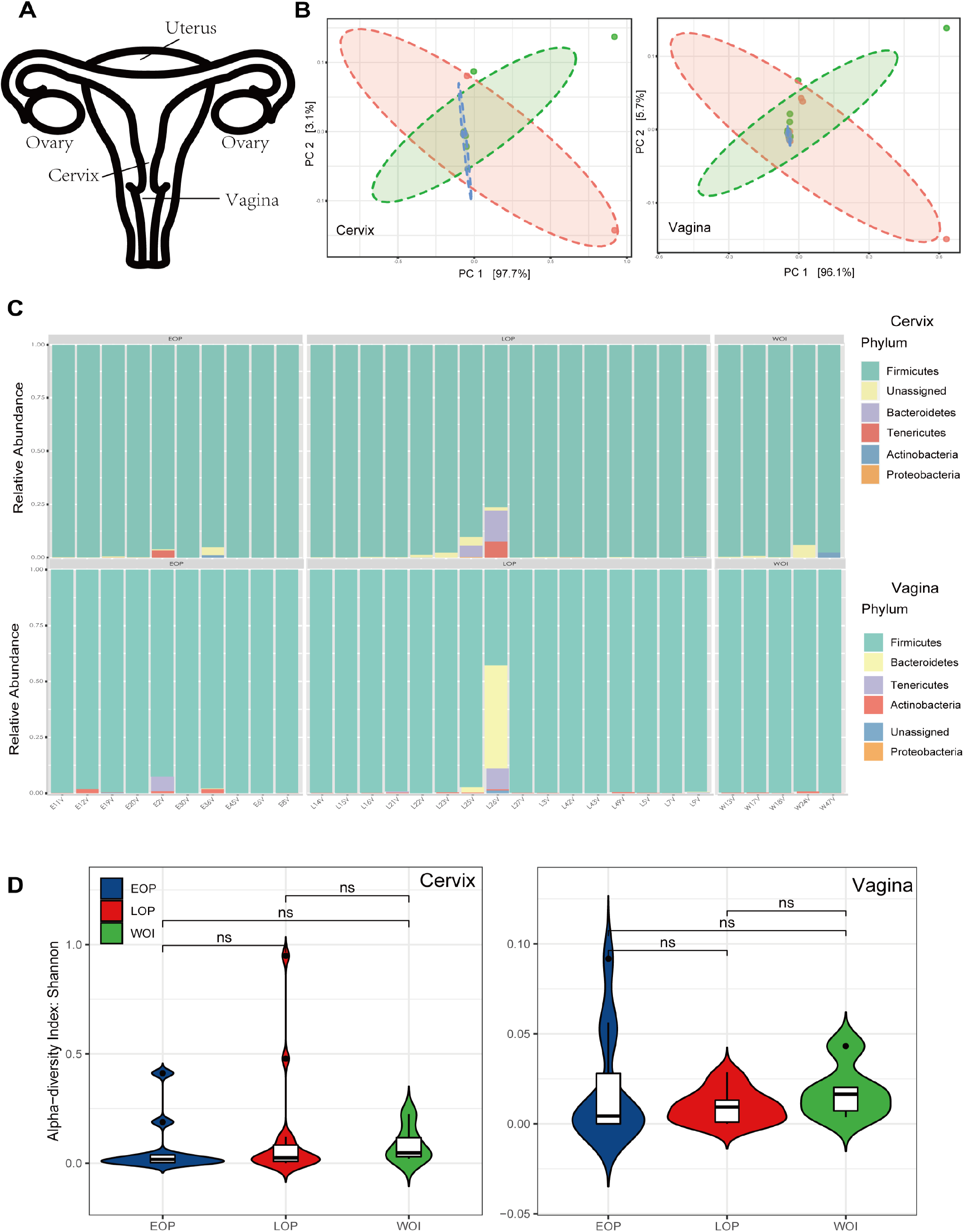
LGT microbiota stays stable over different stages during menstrual interval. A. Structural diagram of the female genital tract. B. Canonical correspondence analysis (CCA) of the LGT (vagina and cervix) microbiota of healthy women at different phases of a menstrual cycle. C. OTU compositions of the LGT (vagina and cervix) microbiota of healthy women at different phases of a menstrual cycle. D. Alpha diversity of the LGT (vagina and cervix) microbiota of healthy women at different phases of a menstrual cycle. LGT: lower genital tract, OTU: operational taxonomic unit

**Figure 2.**
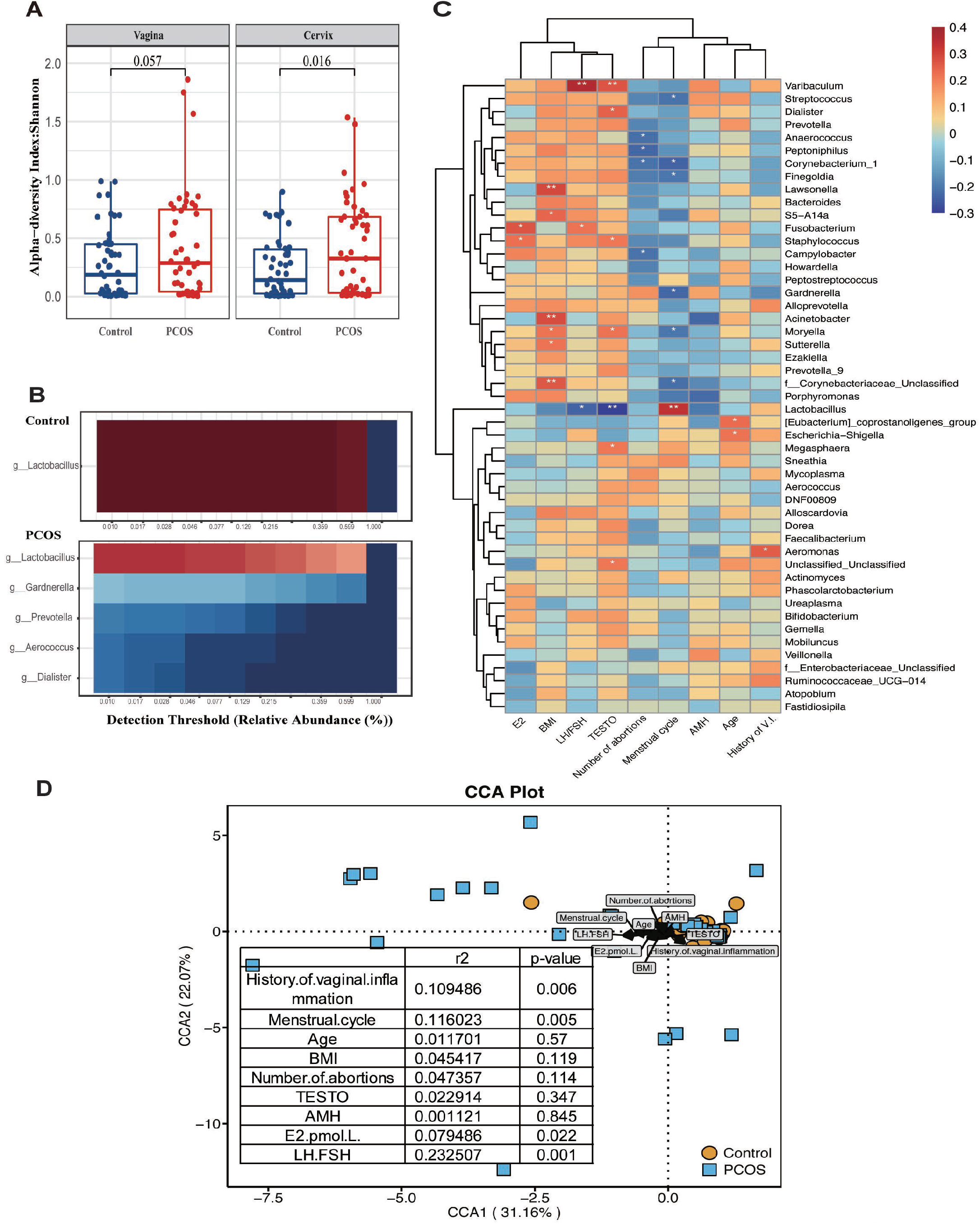
LH/FSH ratio and menstrual cycle frequency are highly relevant to the stability of the LGT microbiota in PCOS patients. A. Comparison of the alpha diversity of the LGT microbiota (vagina (p=0.057) and cervix (p=0.016)) between the PCOS (n=47) and control groups (n=50). B. Heat map of the core microbiome in vaginal samples obtained from the PCOS and control groups. (The core microbiome of the control group is displayed by a heat map that identifies g_*Lactobacillus*, and the core microbiome of the PCOS group is displayed by a heat map that identifies g_*Lactobacillus*, g_*Gardnerella*, g_*Prevotella*, g_*Aerococcus*, and g_*Dialister* as the most prevalent genera). C. Spearman correlation analysis of vaginal bacterial and clinical indexes: red indicates positive correlation; blue indicates negative correlation; *indicates significance, p<0.05; and **indicates significance, p<0.01. D. Canonical correspondence analysis (CCA) of vaginal microbiota composition with clinical indexes (PCOS patients are indicated by blue squares and healthy women by yellow circles. The length of the arrow represents the influence of the factor. If the angle between the arrow and the point is less than 90°, it represents positive correlation; if greater than 90°, it represents positive correlation). LGT: lower genital tract, PCOS: polycystic ovarian syndrome, CCA: canonical correspondence analysis

## Results

### LGT microbiota stays relative stable despite different phases of menstrual cycle

Before study begin, we collected 62 specimens (31 vaginal and 31 cervical swabs) from 22 reproductive-aged healthy women who had regular menstrual cycles (Supplementary Table 1). As hormones and epithelial cells status changed dynamically during a menstrual cycle, those specimens were collected at three different phases of menstrual cycle: (a) follicular phase (within 3 days after the end of menstruation), (b) ovulation phase (1 day before or after ovulation), and (c) planting window phase (7-9 days after ovulation). By 16SrRNA sequencing of collected samples, we found that the LGT microbiota composition stays relative stable at different phases of menstrual cycle, *Lactobacillus* was the most abundant genus in both the vagina and cervical throughout these phases (Figure1B-C). Shannon index showed that the alpha-diversity of vagina and cervical community were not significantly changed between menarche phase, ovulation phase and planting window phase (Figure 1D), and the composition of microorganisms in the vagina and cervical is almost identical, as we previously reported (19).

These results demonstrate that the LGT microbiota of healthy reproductive-aged women stays relatively stable throughout menstrual cycle. Therefore, in the following experiments, we did not limit the sampling of vaginal swabs to a specific period, only avoided the menstrual period. However, as we previously reported, the LGT microbiota is significantly altered in women with PCOS when compared to healthy controls, and the potential mechanism underlying this alteration is still unknown.

### LH/FSH ratio and menstrual cycle frequency are highly relevant to the stability of the LGT microbiota in PCOS patients

Next, we measured clinical indexes (Supplementary Table 2) in a study group of 47 PCOS women and 50 reproductive-aged healthy women, we analyzed the potential relationship between LGT microbiota and clinical data of these women: frequency of menstrual, body mass index (BMI), serum Testosterone level, LH/FSH et al. Shannon index showed that the alpha diversity of the LGT microbiome was significantly increased in PCOS patients when compared to healthy women (Figure 2A): *Lactobacillus* is the most predominant genus in the vagina of healthy women, whereas several other genera (e.g., *Gardnerella* and *Prevotella*) were more abundant in the vagina of women with PCOS (Figure 2B), as same as in the cervical (paper not shown).

Using Spearman correlation analysis, we analyzed the correlation between the LGT flora and clinical indexes of the 97 women (Figure 2C), as microbiological analysis of the vagina and cervical are identical, so we only present the vaginal microbiota analysis results here. We found that the relative abundance of *Lactobacillus spp.* had a positive correlation with menstrual cycle frequency (p<0.01) and negative correlation with serum testosterone level (p<0.01) and LH/FSH ratio (p<0.05). The presence of bacteria genera such as *Gardnerella*, *Finegodia*, *Dialister*, *Acinetobacter*, and *Escherichia-Shigella* were strongly related to reduced menstrual cycle frequency, elevated testosterone level and BMI, and older age. Canonical correspondence analysis (CCA) showed that, among several indexes such as age, body mass index, and sex hormones, LH/FSH ratio was the most relevant factor that was related to the dominant genera in women with PCOS (r^2^=0.233, p=0.001), and menstrual cycle frequency was also strongly related to the LGT organisms (r^2^=0.116, p=0.005) (Figure 2D). CCA analysis revealed that the LH/FSH ratio (Fig.2d, r^2^=0.233, p=0.001) and menstrual cycle frequency (Fig.2d, r^2^=0.116, p=0.005) are most strongly associated with the composition of the microbiota. Considering the negative correlation between LH/FSH ratio and the Lactobacillus genus, along with the positive correlation between menstrual cycle frequency and the Lactobacillus genus (Fig. 2c,2d), we speculated that LH/FSH ratio and menstrual cycle frequency are highly correlated with the stability of the female reproductive tract microbiota of PCOS women, as PCOS women often have a prolonged menstrual frequency and elevated LH/FSH ratio.

### LGT microbiota composition is associated with FET outcomes in PCOS patients

As the LGT microbiota of PCOS patients is dramatically altered compared to that of healthy women, we investigated whether poor reproductive outcomes in women with PCOS (Figure 3A) (Table 1) were also related to the LGT microbiota composition. Here we recruited 72 PCOS patients who were underwent IVF-FET therapy. We sampled LGT swabs (from the vagina and cervical) on the FET day (before transplantation) of those patients, and follow-up visits were conducted and recorded at 14, 28, and 280 days after the FET day. Implantation success was defined as human chorionic gonadotropin ≥5 mg/ml on the 14th day, clinical pregnancy was defined as a visible gestational sac on ultrasound on the 28th day, and live birth was defined as healthy newborn by the 280th day (Figure 3A, 3C),the success rate of clinical pregnancy and live birth in PNB group(PCOS with relative abundance of *Lactobacillus* ≥50%, n=57) was significant higher than PDB group(PCOS patients with relative abundance of *Lactobacillus*<50%, n=15). By mantel test, we found that the genus of *Prevotella* were the most relevant flora that related to clinical pregnancy and live birth (Figure 3B). Also, we found that *Gardnerella* was significantly reduced in the live birth group (p=0.03) (Figure 4C), while *Lactobacillus* was reduced in the implantation failure group at a weak statistical significance (p=0.08) (Figures 4A) (Supplementary Figure 2-4).

**Table 1.** Differences in FET outcomes between the PDB and PNB groups FET: frozen embryo transfer, PDB, PNB

| Group |  | PDB 50%<br>n=15 | PNB 50%<br>n=57 | PCOS total<br>n=72 | Con<br>n=19 |
| --- | --- | --- | --- | --- | --- |
| Implantation Rate |  | 0.33 (n=5) | 0.54 (n=31)* | 0.50 (n=36) | 0.47 (n=9) |
| Clinical pregnancy Rate |  | 0.27 (n=4) | 0.46 (n=26)* | 0.42 (n=30) | 0.42 (n=8) |
| Live birth Rate |  | 0.13 (n=2) | 0.33 (n=19)* | 0.29 (n=21) | 0.37 (n=7) |
| Abortion Rate |  | 0.60 (n=3) | 0.39 (n=12)* | 0.42 (n=15) | 0.22 (n=2) |
| Preterm birth |  | 0 | (n=2) | (n=2) | (n=1) |
| Multiple pregnancy |  | 0 | (n=2) | (n=2) | (n=1) |
| Mode of Delivery | Vaginal | 0.50 (n=1) | 0.21 (n=4) | 0.24 (n=5) | 0.29 (n=2) |
|  | Cesarean | 0.50 (n=1) | 0.79 (n=15) | 0.76 (n=16) | 0.71 (n=5) |
| Age |  | 30.4±2.8 | 29.5±3.4 | 29.8±3.2 | 31.3±3.4 |

#Among the abortion population, there were 2 cases of late abortion, 1 case of cervical insufficiency, and 1 case of induced labor due to fetal malformation
P-value was calculated by $\chi^2$ test, \* $p < 0.05$

**Figure 3.**
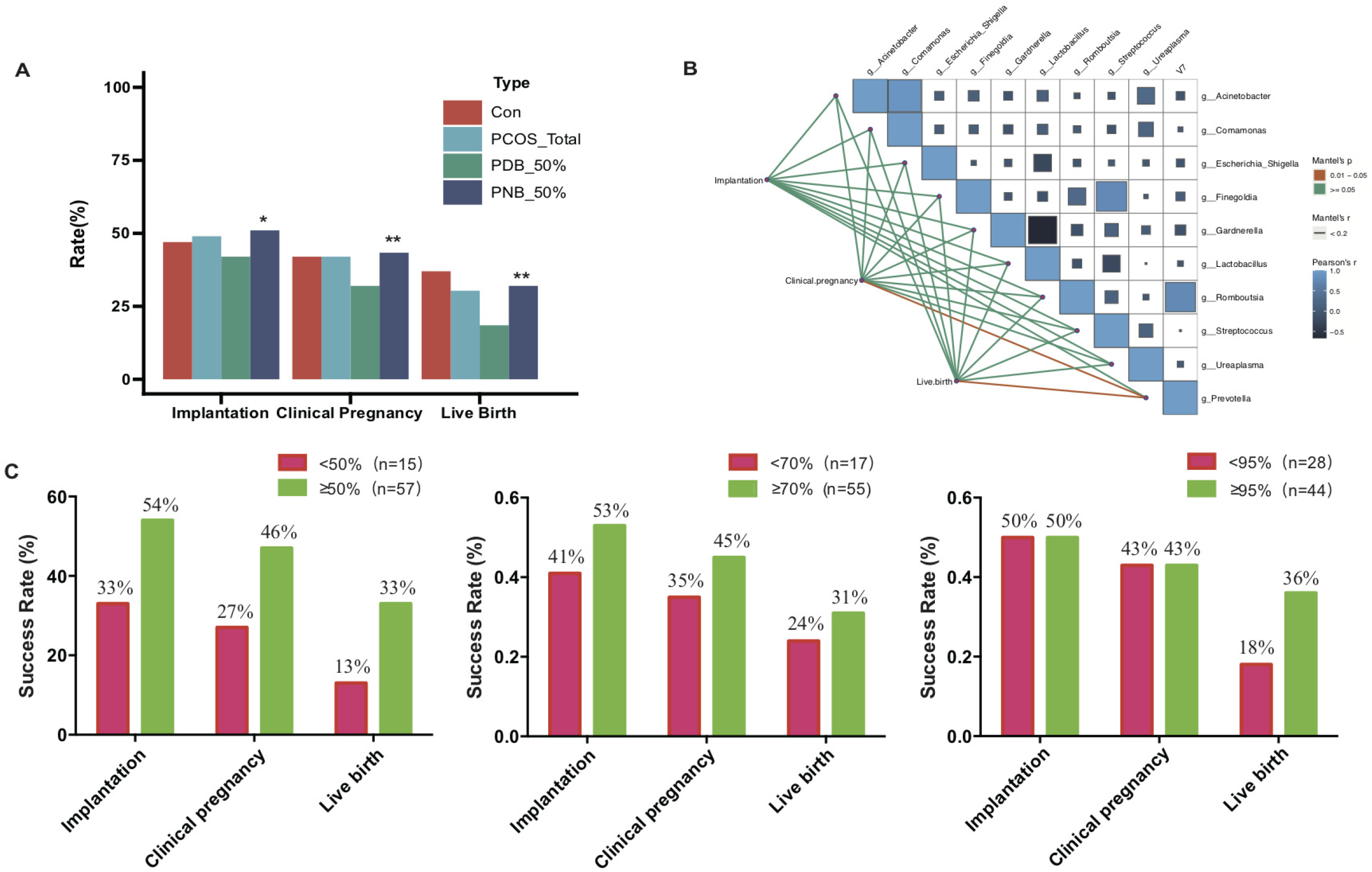
LGT microbiota composition affects the FET outcomes of PCOS patients. A. Comparison of FET outcome among women in the control, PCOS, PDB, and PNB groups who were undergoing FET. B. Correlation coefficients of genus and FET outcome were calculated using Mantel test, and the colored line indicates significant (red) correlations. C. Implantation success, clinical pregnancy, and live birth rates in the PDB (PCOS patients with relative abundance of *Lactobacillus*<50%, n=4) and PNB (PCOS with relative abundance of *Lactobacillus* ≥50%, n=5) groups, consisting of 72 women with PCOS who were undergoing FET. FET: frozen embryo transfer, PCOS: polycystic ovarian syndrome

**Figure 4.**
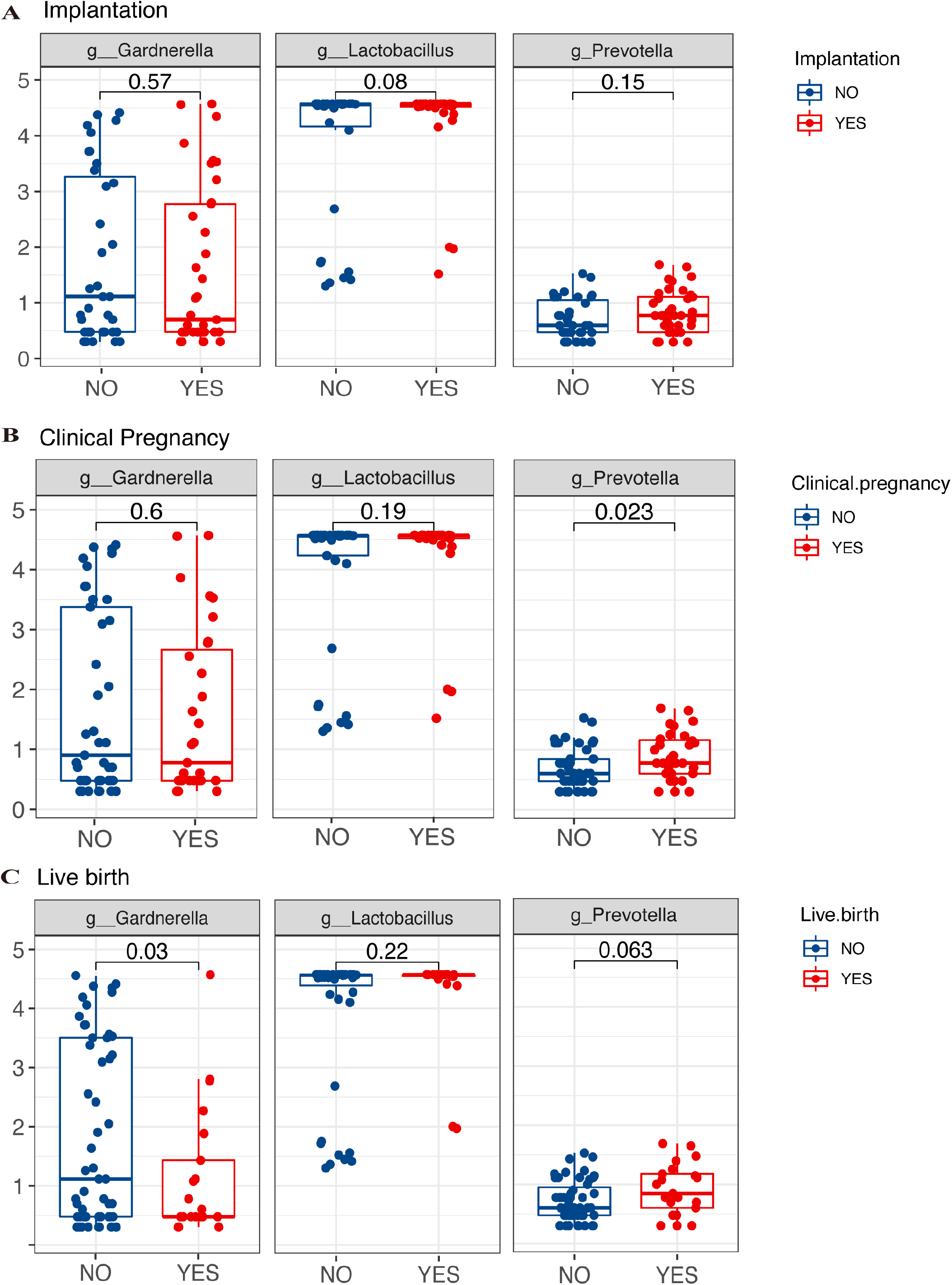
LGT flora were associated with FET outcomes of PCOS women. Comparison of relative abundance of ***Gardnerella spp., Lactobacillus spp., Prevotella spp.*** (the top3 genus abundant in PCOS patients’ LGT) between different pregnancy outcomes as Implantation(A), Clinical pregnancy (B), Live birth(C).

### Alteration of LGT microbiota is related to changes in cervical epithelial cell function in PCOS patients

Further, we doubted whether the function of mucosal epithelial cells interacting with LGT microorganisms is undergo corresponding changes in PCOS women who have an altered LGT microbiota. We randomly selected nine reproductive-aged PCOS patients (approximately half of them had altered LGT microbiota: *Lactobacillus spp.* ≤50%, n=5) and simultaneously collected both LGT swabs and cervical epithelial cells from them. By synchronization analysis of RNA sequencing of the local epithelial cells and 16S rRNA sequencing of the microbes, we found that the gene expression profiles of the cells significantly differed between the PDB (PCOS patients with relative abundance of *Lactobacillus*<50%, n=4) and PNB (PCOS with relative abundance of *Lactobacillus* ≥50%, n=5) groups. The expressions of 115 genes were significantly changed (p<0.05), GO enrichment analyses showed that those genes were mainly involved in biological processes: peptide cross-linking, antimicrobial humoral response, keratinocyte differentiation, cornification, antimicrobial humoral immune response mediated by antimicrobial peptide, etc. (Figure 5A).

**Figure 5.**
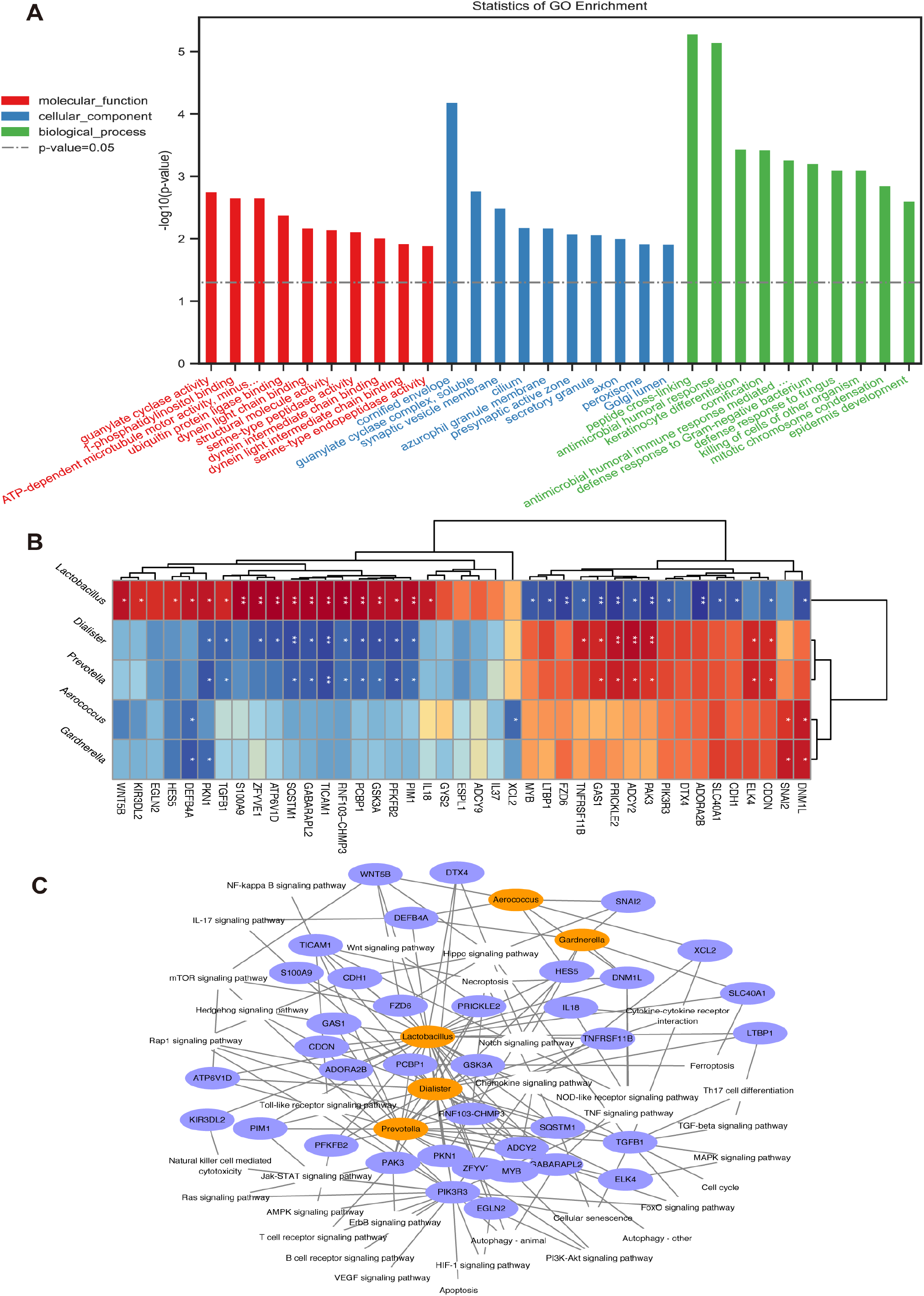
Alteration of the LGT microbiota is related to changes in cervical epithelial cell function in PCOS patients. A. GO enrichment analysis of 497 differentially expressed genes of the cervical epithelial cells between the PDB (PCOS patients with relative abundance of *Lactobacillus spp.* <50%, n=4) and PNB (PCOS patients with relative abundance of *Lactobacillus spp.* ≥50%, n=5) groups. B. The differentially expressed genes that were related to the screened signaling pathways were selected, and correlation analysis was carried out with five target floras: red indicates positive correlation; blue indicates negative correlation; *indicates significance, p<0.05; and **indicates significance, p<0.01. C. The relationship pairs were screened with absolute value of correlation coefficient >0.6, and the gene, flora, and pathway network diagram was drawn: orange indicates the flora, blue indicates the target gene, connection between the gene and flora indicates that there is an expression correlation between the two (that is, the relationship pair has absolute value of correlation coefficient >0.6), and connection between the gene and pathway indicates that the gene is involved in the pathway. LGT: lower genital tract, PCOS: polycystic ovarian syndrome, GO: Gene Ontology

We further mapped the association network between immune response, apoptosis, autophagy, and cell membrane integrity, and five major cervical bacterial genera in patients with PCOS (Figure 5B and 5C, Supplementary Figure 5). Expression of genes that are involved in Toll-like receptor signaling pathway (e.g., *TICAM1*, *PIM1*, *PIK3R3*, and *ATP6VID*) was significantly associated with relative abundance of genera like *Lactobacillus*, *Dialister*, and *Prevotella*. These results indicate that the function of local mucosal epithelial cells has also changed with the flora alternation.

## Discussion

Whether the composition of reproductive tract microorganism stays stable at different stages of the menstrual cycle has been controversial (23, 24), as the composition of microorganisms in LGT will be more unstable due to the continuous outflow of blood and endometrium during menstrual period, so the sampling time of our study completely avoided this period. We selected the sampling time at three different phases of menstrual cycle according to the fluctuate of hormone levels: (a) follicular phase (within 3 days after the end of menstruation), (b) ovulation phase (1 day before or after ovulation), and (c) planting window phase (7-9 days after ovulation). However, we found that different phases of the menstrual cycle (except blooding period) did not affect the composition of the LGT (vaginal and cervical) microbiota in healthy reproductive-aged women although the sex hormone (E_2_, P, LH) levels have changed dramatically at different stages, this result indicated that healthy women who have regular menstrual cycles largely have a stable LGT microbiota composition. As women with PCOS often have prolonged menstrual cycle, whether this abnormality is related to the alternation of LGT microorganisms remains to be studied.

As our previous study demonstrated that the LGT microbiota of women with PCOS is significantly altered when compared to that of healthy controls (19): approximately 28% of women with PCOS have a significantly reduced level of *Lactobacillus spp.* (relative abundance < 50%) in the LGT; however, factors that are responsible for this difference remain unclear. Here we analyzed the correlation between clinical data (such as frequency of menstrual, body mass index (BMI), serum Testosterone level, LH/FSH et al) and LGT microbiota composition in 97 reproductive-aged women (containing 47 PCOS patients and 50 control reproductive-aged women). CCA analysis showed that LH/FSH ratio and menstrual cycle frequency were the most significant clinical indexes that were related to LGT microbiota composition; Spearman correlation analysis also showed that the relative abundance of *Lactobacillus spp.* was mostly associated with menstrual cycle frequency, testosterone level, and LH/FSH ratio. As an increased level of luteinizing hormone (LH) and a decreased level of follicle-stimulating hormone (FSH), is always consistent with elevated levels of insulin, androgen, and anti-Müllerian hormone, which may lead to disorders in the regulation of the menstrual cycle, commonly seen in PCOS women. These factors may collectively promote changes in vaginal microbiota in PCOS patients, but we cannot yet determine the most critical factor, although CCA analysis suggests that menstrual cycle frequency and LH/FSH ratio is the most significantly associated factor. In previously studies, elevated LH/FSH ratio is related to increase in c-reactive protein and free cholesterol, which suggests that increased LH/FSH ratio may accompanied by abnormality of the immune and metabolic status of the body (25, 26).There have no study focused on whether long-term absence of menstruation affects the genital tract microbial composition of reproductive-aged women, we already know that the incidence of vaginitis in postmenopausal women increases, but the cause has often been attributed to low estrogen levels in their body, for women of childbearing age, the relationship between changes in menstrual frequency and reproductive tract microbiota still needs further exploration. However, these results also remind us that for premenopausal women with PCOS, especially those of childbearing age, maintaining a regular menstrual cycle can not only prevent endometrial lesions, but may also be crucial for the homeostasis of their reproductive tract microenvironment, and a better reproductive outcome.

Numerous studies have demonstrated that the microbes live in the genital tract are significantly associated with pregnancy outcomes of reproductive-aged women (5, 27–29), and they may also influence offspring’s health in animal models (30–32). *Lactobacillus spp.* is typically predominant in the LGT of reproductive-aged women (33, 34), in our study, we found that PCOS patients with relatively higher abundance of *Lactobacillus* (>50%) had better FET outcomes: better implantation rates (53%), clinical pregnancy rates (44%), and live birth rates (32%) (Table 1), while patients with relative lower abundance of *Lactobacillus* (≤50%) in LGT had significantly lower rates of implantation (38%), clinical pregnancy (31%), and live birth (13%). We also grouped based on the relatively abundance of *Lactobacillus spp.* for 70% and 90%, however the results didn’t show statistical significance (paper not shown). Therefore, we speculate that the *Lactobacillus spp.* in LGT with relative abundance less than 50% might have a more significant effect on reproductive system function. Next when we grouped and analyzed based on different pregnancy outcomes, we found that the relative abundance of *Gardnerella spp.* significantly lowered in the group of patients who had live births (p=0.03); while other bacteria (means bacteria besides *Lactobacillus spp.*) in live birth group was also lower than the group who were failed to have live birth, despite the lack of statistical significance (p=0.056) (Supplementary Figure 1). These results collectively indicates that the LGT microbiota composition is significantly related with the FET outcome of PCOS women, PCOS women who have a *Lactobacillus spp.* abundant community (especially relative abundance >50%) in LGT might have better pregnancy outcomes. Here we also posed a bold question: Can we improve the reproductive outcomes of PCOS patients by intervening the lower reproductive tract microbiota? Previously research showed that vaginal administration of *Lactobacillus crispatus* in women who had vaginosis resulted in a sustained reduction in genital inflammation and a biomarker of epithelial integrity in vaginal fluid (35), yet whether the cell function of local epithelial changed synchronized remains unclear. Several bacterial species found in *Lactobacillus*-deficient cervicovaginal microbiomes are related to disturbance of immune homeostasis (36, 37), and the relative abundance of *Lactobacillus* in the genital tract is associated with the integrity and function of cervicovaginal mucus (38–43). Interestingly, in our study, we found that the expression level of numerous genes differed between PDB (PCOS patients with relative abundance of *Lactobacillus* < 50%, n=4) and PNB (PCOS with relative abundance of *Lactobacillus* ≥50%, n=5) group, genes such as TICAM3, PKN1, TGFB1, GSK3A, SQSTM1 were positively associated with the abundance of *Lactobacillus spp.*, while PAK3, PRICKLE2, GAS1 were negatively associated. Pathways enriched by those differentially expressed genes mainly involved in pathways such as antimicrobial humoral response, defense response to fungus, killing of cells of other organism, et al, indicates that PCOS women who with a relative lower abundance of *Lactobacillus* in LGT may highly accompanied by an immune-imbalance environment in the LGT. Besides, pathways such as keratinocyte differentiation, cornification were also changed dramatically between those two groups, as change in epidermal keratinization can affect the formation of the skin barrier; therefore, disorder of the LGT flora in women with PCOS may be related to disorder of the local skin mucosal barrier system and to local inflammatory reactions. These results indicates that the LGT microbiota of women with PCOS may play an important role in the local mucosal epithelial barrier and immune homeostasis: PCOS women with relative abundance of *Lactobacillus spp.* ≥50% in the LGT are more likely to have a better epithelial function and better FET outcomes, and we need more clinical research to verify whether the vaginal *Lactobacillus spp.* complication can improve the fertility outcomes of PCOS women.

## Conclusions

Poor reproductive outcomes have always been an important issue that troubles PCOS patients of reproductive age. In the past, ovulation disorders and poor egg quality were often considered as the main reasons. However, through ovulation induction therapy and even IVF assisted reproduction, a considerable number of PCOS patients still face embryo implantation failure, high miscarriage rates and premature birth rates; we considered that the role of genital tract microenvironment in this issue had been underestimated. In this study, through three independent study groups, we revealed that the LGT microbiota is closely related with reproductive system function (clinical indexes such as LH/FSH, menstrual frequency; and cervical cell function) and IVF-FET outcome in PCOS women, besides PCOS women with relative abundance of *Lactobacillus spp*. <50% in LGT is more likely to have a poorer FET outcome and dysfunction in cervical cell. All these results indicated the critical role of *Lactobacillus spp*. in PCOS women’s reproductive system, further research might need to verify the importance of *Lactobacillus spp*. by vaginal intervention therapy, and more research needed to focus on the specific mechanisms of interaction between LGT microbiota and reproductive system in PCOS patients, to provide novel ideas and angles for clinical diagnosis and treatment.

## Authors’ roles

Yaoyao Tu contributed to sample and data collection and analysis, wrote the first draft, and revised all versions of the manuscript. Yixiang Zhang performed data analysis. Huixi Chen, and Bin Weicontributed to the sample and data collection. Guolian Ding, Xi Dong and Jianzhong Sheng supervised data interpretation and manuscript writing. Hefeng Huang was the principal investigator in this research. Hefeng Huang and Li Jin had primary responsibility for the final content and supervised and contributed to all aspects of the study. All authors read and approved the final manuscript.

## Acknowledgements

We thank all the women who participated in this study and the researchers for their efforts.

## Data Availability

All sequence data are available at the National Center for Biotechnology Information. The 16S rRNA sequences were deposited under accession numbers SAMN21619565-21619724, SAMN21619392-21619411, SAMN21619461-21619544, SAMN21619730-21619738. RNA-Seq data were deposited under BioProject ID PRJNA766136, SRA accession number SUB10427247-10428494.

## Supplementary data

**Supplementary Table 1.** Patients list in study group 1

EOP: basic endocrinology stage, LOP: ovulation stage, WOI: implantation stage

**Supplementary Table 2.** Clinical indexes of 47 women with PCOS and 50 reproductive-aged healthy women PCOS: polycystic ovarian syndrome

**Supplementary Figure 1.** Grouping by different FET outcomes: implantation, clinical pregnancy, and live birth, compared with the relative abundance of *Lactobacillus spp.* and other microbes (means microbes except *Lactobacillus spp.*)

FET: frozen embryo transfer

**Supplementary Figure 2.** Grouping by implantation success or failure, the relative abundance of top 10 bacteria that lives in LGT.

**Supplementary Figure 3.** Grouping by clinical pregnancy, the relative abundance of top 10 bacteria that lives in LGT.

**Supplementary Figure 4.** Grouping by live birth, the relative abundance of top 10 bacteria that lives in LGT.

**Supplementary Figure 5.** A. KEGG analysis of 497 differentially expressed genes of cervical epithelial cells between the PDB (PCOS patients with relative abundance of *Lactobacillus spp.* <50%, n=4) and PNB (PCOS patients with relative abundance of *Lactobacillus spp.* ≥50%, n=5) groups. B. Alpha diversity between the PDB and PNB groups.

KEGG: Kyto Encyclopedia of Genes and Genomes

